# On Breathing Variability in the Tree Shrew

**DOI:** 10.64898/2026.08.13.744653

**Authors:** Devin Bishop, Julianna Saxenna, Shahriar SheikhBahaei

## Abstract

Tree shrews (*Tupaia belangeri*) are increasingly used in comparative neuroscience, yet their respiratory physiology remains poorly characterized. We quantified spontaneous breathing and respiratory rhythm variability in awake adult tree shrews (n = 10; 5 males, 5 females) using whole-body plethysmography. Respiratory frequency decreased by approximately 16% with acclimatization to the recording chamber, while respiratory timing, body-mass-normalized respiratory amplitude, inspiratory flow, and minute ventilation remained relatively stable. After acclimatization, mean respiratory parameters were similar between sexes, but short-term breath-to-breath variability (SD1) was greater in males than females, whereas SD2 was comparable. These findings establish baseline respiratory characteristics in awake tree shrews and identify sex-dependent differences in short-term respiratory rhythm stability.

## Introduction

The basic rhythm of breathing is generated continuously by brainstem respiratory networks (1,2); however, respiratory rate, depth, timing, and breath-to-breath variability are dynamically shaped by metabolic demand, chemosensory feedback, behavioral state, and arousal (3–10). Variability in respiratory cycle duration can therefore provide information that is not captured by mean respiratory rate or ventilation alone (11–14). Poincaré analysis, in particular, separates short-term breath-to-breath variability (SD1) from longer-timescale variability (SD2), providing a useful framework for quantifying respiratory rhythm dynamics (13). Previous work has demonstrated that regularity of breathing is shaped by both peripheral sensory feedback from the carotid bodies and central neural mechanisms, including pontine Kölliker–Fuse circuits, brainstem catecholaminergic neurons, serotonergic signaling, and astrocyte-dependent regulation of respiratory networks (8,15–19). More recently, we applied automated breath-by-breath analysis and Poincaré-based measures to awake rats and common marmosets, demonstrating that respiratory pattern and regularity can be quantified reliably in freely behaving rodents and non-human primates (6,14).

Biological sex is an important source of variation in respiratory physiology. Differences between males and females arise from both respiratory-system anatomy and sex-dependent regulation of neural and chemosensory control mechanisms (20). Human studies further suggest that sex can influence the stability and variability of breathing. Premenopausal women are less susceptible than men to hypocapnia-induced apnea during sleep, testosterone administration increases susceptibility to ventilatory instability in women, and inhibition of testosterone conversion increases breathing stability in healthy men (21–23). Recent measurements in healthy young adults aged 18–23 years have also reported greater spontaneous respiratory variability in men than women (24), while studies of chemoreflex control indicate greater ventilatory loop gain in men under hypoxic conditions (25,26). These differences, however, are not universal across mammals. In our previous study of common marmosets, baseline respiratory rate, body-mass-normalized tidal-volume estimates, ventilation, and respiratory irregularity quantified by SD1 and SD2 were comparable between males and females (14). Thus, the extent to which sex-dependent respiratory variability is conserved across mammalian lineages remains unclear.

Tree shrews (*Tupaia belangeri*) are capable of addressing this inquiry. They belong to the order Scandentia within Euarchontoglires and exhibit genomic and neurobiological characteristics that render them particularly valuable for comparative studies involving rodents and primates (27,28). Despite their increasing use in systems and behavioral neuroscience, fundamental features of respiratory physiology in awake tree shrews remain largely undefined. Here, we characterized spontaneous breathing in awake tree shrews using whole-body plethysmography recordings and quantified respiratory rate, respiratory amplitude, inspiratory and expiratory timing, inspiratory flow, ventilation, and breath-to-breath irregularity. We further compared these parameters between male and female animals to determine whether biological sex influences either the average breathing pattern or its temporal stability. By establishing baseline respiratory characteristics in tree shrews, this study provides a physiological framework for their growing use in comparative and systems neuroscience and allows sex-dependent features of respiratory control to be evaluated across rodents, tree shrews, non-human primates, and humans.

## Materials and Methods

### Animals

Adult tree shrews (*Tupaia belangeri*) were used to characterize spontaneous breathing behavior. The final analysis included 10 adult animals (5 males and 5 females; age: 14 ± 3 months; range: 11-20 months). Animals were housed in a temperature-controlled facility under a 12 h light /12 h dark cycle with food and water available *ad libitum*. All procedures were performed in accordance with the National Institutes of Health Guide for the Care and Use of Laboratory Animals and were approved by the Institutional Animal Care and Use Committee of Stony Brook University.

### Measurement of respiratory activity

Respiratory activity was measured non-invasively for 45 minutes using a whole-body plethysmography chamber, adapting procedures previously used to characterize breathing behavior in awake rodents and common marmosets (8,14,16,29). Awake tree shrews were placed individually in a transparent plethysmography chamber (∼3 L) which was flushed with room air (21% O_2_, 79% N_2_), at a rate of 2.5 L min^−1^ and allowed to move freely throughout the recording. The chamber was continuously ventilated with room air during measurements of spontaneous baseline breathing. Recordings were performed under ambient laboratory conditions within a consistent daytime window (10:00–14:00). Pressure fluctuations produced by respiratory movements were recorded continuously and digitized for subsequent offline analysis. Respiratory data were acquired with a Power1401 (CED; RRID: SCR_017282) interface and transferred to Spike2 software (CED; RRID: SCR_000903; **Figure 1A**). For each animal, a baseline recording period, the first and last 10-min windows, was analyzed. Respiratory signals containing major movement artifacts or non-respiratory pressure transients were excluded before breath-by-breath analysis.

**Figure 1.**
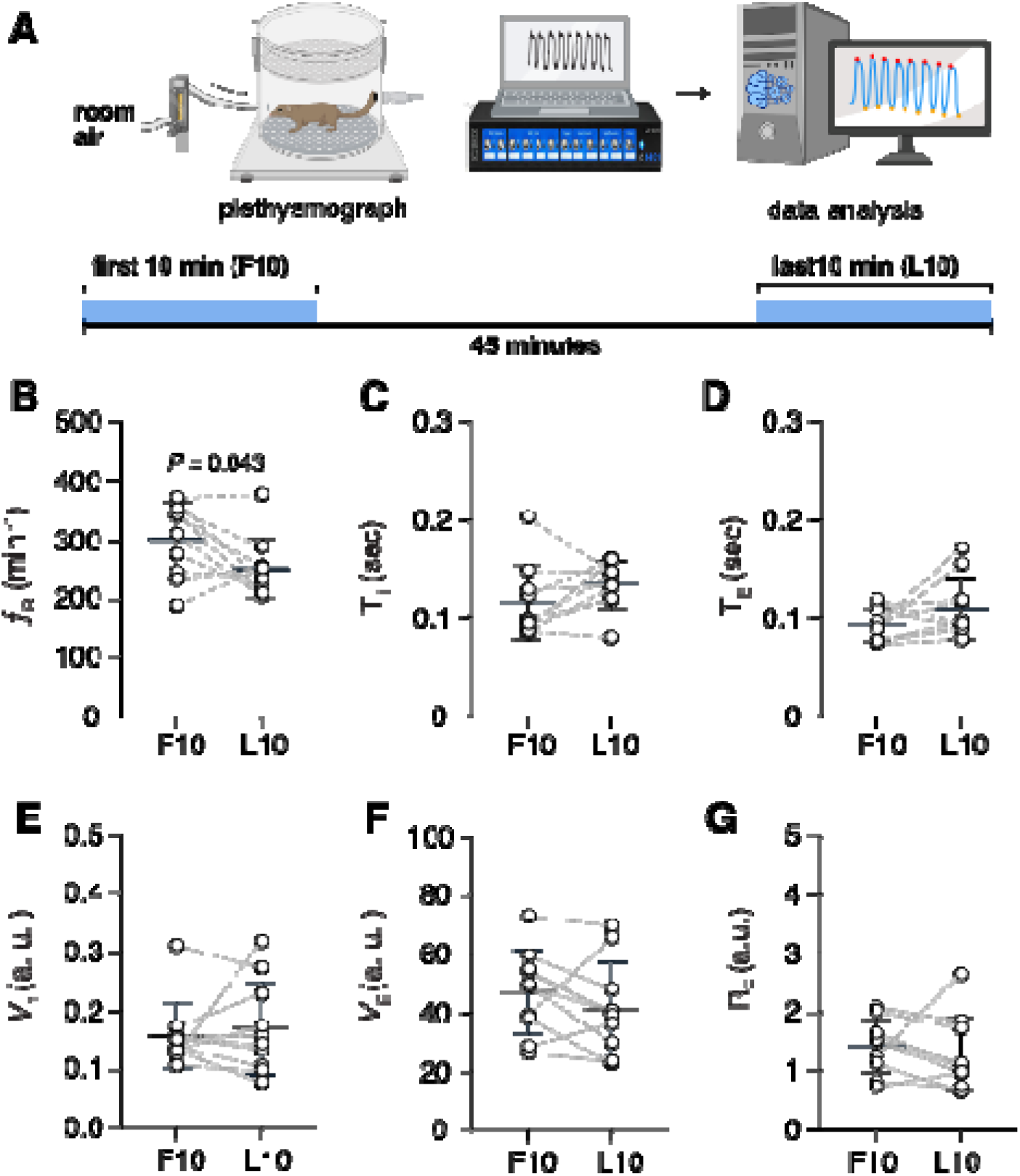
Respiratory frequency decreases during acclimatization to the recording chamber while other respiratory parameters remain relatively stable. (**A**) Schematic of the experimental design. Awake adult tree shrews were recorded in room air for 45 min in a whole-body plethysmography chamber. Respiratory measurements from the first 10 min of chamber exposure (F10) and the last 10 min after acclimatization (L10) were compared. Biorender.com was used to make the schematic. (**B–G**) Paired comparisons of respiratory frequency (*f*_R_; **B**), inspiratory time (T_I_; **C**), expiratory time (T_E_; **D**), body-mass-normalized tidal volume (V_T_; **E**), minute ventilation (V_E_; **F**), and inspiratory flow (R_F_ = V_T_ / T_I_; **G**) between F10 and L10. Each circle represents one animal (n = 10), with measurements from the same animal connected by dashed lines. Horizontal bars and error bars indicate mean ± SD. Statistical comparisons were performed using two-tailed paired *t*-tests. Data without a *P* value were not statistically significant. a.u.: arbitrary unit.

### Respiratory signal analysis

Respiratory signals in the first 10 minutes (F10) and the last 10 minutes (L10) were analyzed offline using a Python-based analysis pipeline adapted from our previously described automated respiratory analysis workflow (14) (**Figure 1A**). Signal processing and breath detection were performed using Python packages including NeuroKit2, NumPy, and Pandas. Individual respiratory cycles were identified from successive respiratory troughs and peaks, and breath-by-breath measurements were extracted from the cleaned signal. For each respiratory cycle, respiratory frequency (f_R_), total respiratory cycle duration (T_TOT_), inspiratory time (T_I_; duration of the inspiratory phase), expiratory time (T_E_; duration of the expiratory phase), and trough-to-peak respiratory amplitude were calculated. Respiratory frequency was expressed as breaths per minute. The T_I_ / T_E_ ratio was calculated for each recording as an additional measure of respiratory timing. The trough-to-peak respiratory amplitude was used as a proxy for tidal volume (V_T_). To account for differences in body size between animals, amplitude-dependent respiratory variables were normalized to body weight. Mean inspiratory flow rate (R_F_) was calculated as the ratio of normalized tidal volume to inspiratory time (V_T_ / T_I_). Minute ventilation (V_E_) was estimated as the product of normalized tidal volume and respiratory frequency: V_E_=V_T_×f_R_

### Respiratory rhythm variability

Breath-to-breath respiratory variability was quantified using Poincaré analysis of respiratory cycle duration, as previously described for rodents and common marmosets (8,14). For each recording, T_TOT_(n) was plotted against the duration of the subsequent respiratory cycle, T_TOT_(n+1). Two measures of respiratory rate variability were derived from the resulting Poincaré distribution as described before (12,14). SD1 represents dispersion perpendicular to the line of identity and therefore quantifies short-timescale, breath-to-breath variability. SD2 represents dispersion along the line of identity and reflects variability occurring over longer timescales. SD1 and SD2 were calculated from the standard deviation of T_TOT_ and the standard deviation of successive differences in T_TOT_: SD_1_ ^2^ = ½ SD _SD_^2^ and SD_2_ ^2^ = 2SDT _TOT_^2^ – ½ SD _SD_^2^, where SD_SD_ is the standard deviation of successive differences in T_TOT_ and SDT_TOT_ is the standard deviation in T_TOT_. Higher SD1 values therefore indicate greater cycle-to-cycle respiratory variability, whereas higher SD2 values indicate greater longer-timescale variation in respiratory cycle duration (14). Effect size was quantified using Hedges’ g, calculated as the standardized difference between group means using the pooled standard deviation and corrected for small-sample bias (30).

### Sex comparisons and statistical analysis

Respiratory parameters were compared between male and female tree shrews. Each animal represented an independent biological replicate, with one baseline respiratory recording contributing to the final analysis. The final dataset consisted of five males and five females. Data were analyzed in Prism 11 (GraphPad, Inc; RRID: SCR_002798) and are presented as mean ± SD. Male and female groups were compared using two-tailed unpaired t-test, Welch’s *t*-test, two-way repeated-measures ANOVA, and Hedges’ g as appropriate.

## Results

### Respiratory rate decreases over the course of recording in awake tree shrews

We first examined whether spontaneous breathing changed over the course of recording by comparing the first and last 10 min of each recording. Across animals (n = 10), respiratory rate (f_R_) decreased from 304 ± 63 breaths /min during the first 10 min (F10) to 254 ± 51 breaths /min during the last 10 min (L10; *P* = 0.043, paired *t*-test; **Figure 1B**).

We next examined this change separately in male and female tree shrews. In females, respiratory rate decreased from 308 ± 48 breaths /min during the first 10 min to 231 ± 19 breaths / min during the last 10 min (*P* = 0.02, paired t-test). In males, respiratory rate (f_R_) decreased from 299 ± 81 to 277 ± 63 breaths /min, although the magnitude of this change was smaller and more variable (P = 0.6, paired t-test). The change in f_R_ over time did not differ between males and females (sex × time interaction, *P* = 0.2).

Other respiratory parameters remained relatively stable over the recording period. Inspiratory time (T_I_) was 0.12 ± 0.04 s during the first 10 min and 0.13 ± 0.02 s during the last 10 min (*P* = 0.13; **Figure 1C**), whereas expiratory time (T_E_) was 0.09 ± 0.02 and 0.11 ± 0.03 s, respectively (*P* = 0.12; **Figure 1D**). The T_I_ / T_E_ ratio was also unchanged (1.2 ± 0.2 vs. 1.3 ± 0.3; *P* = 0.6). Respiratory amplitude (V_T_), normalized to body mass, was similar between the first and last 10 min (0.16 ± 0.06 vs. 0.17 ± 0.08 a.u.; *P* = 0.6; **Figure 1E**), as were minute ventilation (V_E_; 47 ± 14 vs. 42 ± 16 a.u. /kg /min; *P* = 0.2; **Figure 1F**) and inspiratory flow rate (R_F_; 1.4 ± 0.5 vs. 1.3 ± 0.6; *P* = 0.5; **Figure 1G**). These data indicate that awake tree shrews exhibit a time-dependent slowing of respiratory frequency during recording without a corresponding broad change in respiratory amplitude, timing, or ventilation.

### Respiratory parameters after acclimatization are similar between male and female tree shrews

We next characterized respiratory physiology during the last 10 min of recording and compared these parameters between male and female animals. Across all tree shrews, f_R_ during this period was 254 ± 51 breaths /min, T _I_ was 0.13 ± 0.02 s, T_E_ was 0.11 ± 0.03 s, and the T_I_ / T_E_ ratio was 1.3 ± 0.3. V_T_ was 0.17 ± 0.08 a.u., inspiratory flow rate was 1.3 ± 0.6, and V_E_ was 42 ± 16 a.u. Respiratory timing was comparable between sexes. f_R_ was 277 ± 63 breaths /min in males and 231 ± 19 breaths /min in females (*P* = 0.2). T_I_ did not differ between males and females (0.13 ± 0.03 vs. 0.14 ± 0.02 s; *P* = 0.3), nor did T_E_ (0.10 ± 0.03 vs. 0.12 ± 0.03 s; P = 0.4). The T _I_ / T_E_ ratio was virtually identical between males and females (1.3 ± 0.3 vs. 1.3 ± 0.4; *P* = 0.99). Although male animals weigh more than females (200 ± 19 vs. 174 ± 16 g; *P* = 0.047), V_T_ was similar between males and females (0.17 ± 0.08 vs. 0.17 ± 0.09 a.u.; *P* = 0.9). R_F_ (1.3 ± 0.4 vs. 1.3 ± 0.8; *P* = 0.8) and V_E_ (44.8 ± 15.4 vs. 38.8 ± 17.5 a.u.; *P* = 0.6) were likewise comparable between sexes.

### Short-term respiratory variability differs between male and female tree shrews

We next examined respiratory rhythm variability during the final 10 min. SD1, which reflects short-term breath-to-breath variability, was higher in male than female tree shrews (61 ± 10 vs. 38 ± 14 a.u. ; *P* = 0.023; **Figure 2A**). The magnitude of this sex difference was large (Hedges’ g = 1.64). In contrast, SD2, which reflects variability over longer timescales, was similar between males and females (101 ± 28 vs. 102 ± 36 a.u.; *P* = 0.9; **Figure 2B**). Thus, although mean respiratory timing, amplitude, drive, and ventilation were similar between sexes, male tree shrews exhibited greater short-timescale respiratory variability. These findings identify a sex-dependent difference in breath-to-breath respiratory dynamics that is selective for SD1 rather than a generalized increase in respiratory variability.

**Figure 2.**
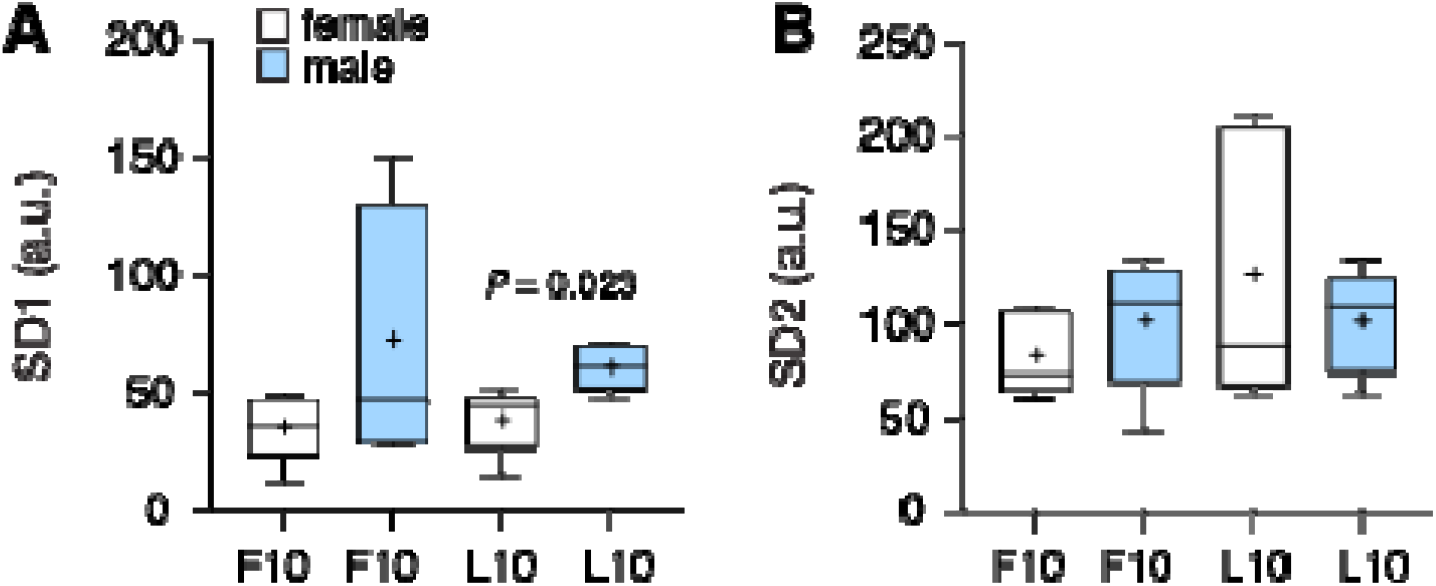
Sex differences in respiratory rhythm variability in tree shrews. Respiratory rhythm variability was quantified from total respiratory cycle duration (TTOT) using Poincaré analysis during the first 10 min of chamber exposure (F10) and the last 10 min after acclimatization (L10). (**A**) Short-term respiratory rhythm variability (SD1) in female and male tree shrews. After acclimatization, SD1 was significantly greater in males than in females (*P* = 0.023, two-tailed Welch’s *t*-test). (**B**) Long-term respiratory rhythm variability (SD2) in females and males during F10 and L10; no significant sex difference was detected. Boxes show the distribution for each group (n = 5 animals / sex); whiskers represent minimum to maximum; horizontal lines indicate medians and + symbols indicate means. a.u.: arbitrary unit.

## Discussion

Here, we characterized spontaneous breathing in awake tree shrews and identified two features of their respiratory physiology. First, respiratory frequency (f_R_) decreased over the course of recording, whereas respiratory timing, amplitude, inspiratory flow, and minute ventilation remained relatively stable. Second, after acclimatization to the recording chamber, mean respiratory parameters were comparable between males and females, but short-term breath-to-breath variability was greater in males. The latter effect was selective for SD1, with essentially identical SD2 values between sexes. Together, these findings indicate that respiratory output in awake, freely moving tree shrews is influenced by arousal state and biological sex, with sex differences being expressed primarily in the temporal stability of breathing rather than in mean ventilatory parameters.

Respiratory frequency after acclimatization to the recording chamber averaged approximately 254 breaths /min, substantially higher than previous values reported for tree shrews. (31) reported a resting respiratory frequency of approximately 97 breaths /min in *Tupaia glis*, whereas (32) reported approximately 120 breaths /min in *T. belangeri* before anesthesia. These measurements, however, were obtained using visual assessment of respiratory movements rather than continuous breath-by-breath plethysmography. The higher respiratory frequency observed upon exposure to the recording chamber may reflect mild arousal or stress associated with the novel environment. As the animals acclimatized to the chamber, respiratory frequency decreased by approximately 16% (from 304 to 254 breaths /min), while respiratory amplitude, timing, and ventilation remained relatively stable. This decline was particularly apparent in females (a decrease of ∼ 25%), although the effect of recording time did not differ between sexes. This pattern is consistent with a reduction in novelty- or arousal-associated respiratory drive rather than a broad change in ventilatory demand. In fact, breathing is strongly modulated by arousal, sensory input, locomotor state, and ongoing behavior in addition to metabolic and chemosensory requirements (3,4,8,10,33). Thus, respiratory measurements obtained shortly after placement in a recording chamber may not represent the same physiological state as measurements obtained later in the session. This temporal dependence should be considered when establishing baseline respiratory values in tree shrews and when comparing respiratory measurements across experimental paradigms.

Despite clear differences in body mass, males and females showed similar respiratory frequency, inspiratory and expiratory timing, respiratory amplitude, inspiratory flow, and minute ventilation after acclimatization to the recording chamber. The principal sex-dependent feature was instead respiratory rhythm variability. SD1 was approximately 60% higher in males than in females and showed a large standardized effect size, whereas SD2 did not differ between sexes. Because SD1 reflects rapid cycle-to-cycle fluctuations, while SD2 captures variability occurring over longer timescales, this dissociation suggests that sex preferentially influences moment-to-moment regulation of respiratory timing rather than producing a generalized increase in respiratory instability. Respiratory variability can arise from multiple interacting processes, including peripheral chemosensory feedback, pontine regulation of respiratory timing, serotonergic and catecholaminergic modulation, and astrocyte-dependent regulation of respiratory networks (8,15–18). Our data therefore suggest a sex-dependent difference in the moment-to-moment regulation of respiratory rhythm.

The male-associated increase in SD1 in tree shrews parallels recent observations in healthy young adults, in whom spontaneous respiratory variability was greater in men than women (24). In contrast, we previously found comparable SD1 and SD2 values in male and female common marmosets (14), suggesting that sex effects on respiratory variability are not uniformly conserved across mammals. Whether this difference reflects sex-dependent chemoreflex gain, neuromodulatory regulation, hormonal influences, or differential coupling of breathing to behavioral state remains to be determined.

The greater short-term respiratory variability observed in males may also be relevant to vocal motor control, given the close neural and biomechanical coordination of respiratory and laryngeal systems during speech (19,34–36). Speech requires precise coordination of respiratory, laryngeal, and articulatory motor systems, and previous studies have identified atypical interactions between respiratory control systems during speech in some individuals who stutter (34). This relationship is particularly intriguing given the marked male predominance of persistent stuttering and the greater probability of recovery in females (37,38). In parallel, we have found increased respiratory irregularity and altered chemoreflex control in *Gnptab*-mutant mice carrying a mutation associated with stuttering (39,40). Together, these observations suggest that respiratory rhythm stability is one component of a broader motor-control phenotype relevant to vocal fluency. Greater baseline respiratory variability in males could increase the demands placed on the coordination of respiratory and vocal motor networks, although whether this contributes to the sex bias observed in speech disorders remains to be determined.

## Acknowledgements

We thank Sophia Zorakhsh, Wendy Akmentin, and the Stony Brook DLAR staff, including Dr. Rachel Brownlee (Director), Kristin Lasek, Laura Readler, and Kaleigh Spetta, for technical support. We also thank the Max Planck Institute Florida (Amanda Coldwell, Susan Freling, Nicole Shultz, and members of the Fitzpatrick Lab) for technical support in establishing the colony.

